# Shotgun metagenomic analysis of the skin mucus bacteriome of the common carp (*Cyprinus carpio*)

**DOI:** 10.1101/2023.06.12.544578

**Authors:** Márton Papp, Adrienn Gréta Tóth, Sára Ágnes Nagy, Károly Erdélyi, Gergely Maróti, Niamh Cox, László Czikk, Máté Katics, László Békési, Norbert Solymosi

## Abstract

The skin mucus bacteriome of fish plays an important role in the health of their hosts. Despite the economic importance of the common carp (*Cyprinus carpio*), research on its skin bacteriome composition is still missing. To date, most studies on the composition of fish skin bacteriome have used amplicon sequencing, despite the limitations associated with this method. In our study, a shotgun metagenomic approach was applied to characterize the external mucus bacteriome of 8 carp specimens from two different ponds on a fish farm in Hungary. Besides the carp samples, water was also sequenced from the two corresponding ponds. Each carp skin sample was dominated by the phylum *Proteobacteria*, followed by *Actinobacteria, Bacteroidota, Firmicutes, Cyanobacteria* and *Planctomycetota*. Additionally, we have found strong concordance between the water and carp skin mucus samples, despite most studies describing an opposite relationship. Furthermore, shotgun metagenomics allowed us to apply functional annotation to the metagenomes, which revealed several metabolic functions. We present, to our knowledge, the first description of the common carp (*Cyprinus carpio*) skin mucus bacteriome. Even though our results showed a high level of host genome contamination, we could still provide valuable insight into the external bacterial community of this species. The presented data can provide a basis for future metagenome studies of carp or other fish species.

## Introduction

The external mucus of fishes provides a home to a wide variety of microorganisms, as it is known for other species^1,2^. These microorganisms, however, are not hitchhikers of their hosts but possess important functional roles, such as defence against various pathogens^3–5^. The colonization of the skin mucus of fishes is assumed to originate from the surrounding water, which process may even start at the larval stage^3^. However, the fish skin bacteriome composition is influenced by several factors such as stress^1^, water pH level^6^ or other environmental influences^7–9^. Furthermore, even the genetics and diet of the host species can have an effect on its structure^1,8,10,11^. Moreover, even within a single individual, different body parts may show differences in microbiome composition^12^.

The microorganisms that inhabit the skin are important for the well-being of their hosts^3–5^. They might even play a practical role in the maintenance of the health of these animals, for example, as an indicator of various pathological conditions^13,14^, or as a source for potential future probiotics^15^. Due to the economic importance of the common carp among freshwater fish species^16–18^, efforts to protect their health are particularly important. However, it should be noted that studies on the bacteriome and microbiome of this species are underrepresented compared to other species, especially considering the skin mucus bacteriome. For this reason, it would be beneficial to increase our knowledge on the bacteriome of the common carp as well.

Despite the long history of the study of the microbial and bacterial community of the outer surface of fishes^19,20^, it has recently received much more attention due to the advent of next-generation sequencing (NGS) technologies^4,14^. However, it is important to note that 16S rRNA gene-based methods have been used in the majority of such studies on the bacteriome of fish skin mucus^4,14,21^. A review article from 2021 listed only one paper using shotgun metagenomics for the analysis of the external surface of eels^21,22^. Beyond which, to the best of our knowledge, we are aware of only one further shotgun metagenomics study from 2020^23^ investigating the fish skin metagenome of cartilaginous and bony fishes from an evolutionary perspective. Despite the conflicting results on the effectiveness of the two methods in revealing microbial community structure^24–27^, it is certain that shotgun sequencing-based methods have the major advantage of providing much greater insight into the functional organization of microbial communities^14,24,25^.

Our study aimed to investigate the composition of the bacteriota inhabiting the carp’s external mucus using shotgun metagenomics. Our results may thus provide a basis for future studies^28^ either using similar methodology or for comparison with 16S rRNA gene-based ones. In addition, the use of shotgun metagenomics for studying the skin mucus bacteriome of the common carp allowed us to analyze the functional and metabolic potential associated with it.

## Materials and Methods

### Samples

All carp (*Cyprinus carpio*) skin mucus samples were taken from two separate ponds at a fish farm in Hungary. Sampling was performed on 15.09.2021. The skin mucus was collected in sterile tubes with plastic tools that were changed between each subject. Mucus collection was restricted to the lateral line region of the fishes so faecal matter wouldn’t impair our results. Furthermore, great care was taken during handling to avoid contamination of the samples. Four samples were taken at each pond. Metadata on samples can be found in Supplementary File 1.

At the farm where samples were collected, both scaly and mirror carp phenotypes are kept. During the sample collection, we could sample two of each at one pond, however, only one scaly and three mirror carp at the other. Furthermore, it is worth mentioning that two specimens from pond 1 had ulcers on their skin, otherwise, all sampled fish appeared to be healthy. Details on the metadata on each sample, along with the number of reads used for classification, can be found in Supplementary File 1.

In addition to the skin mucus samples, water was collected from each pond. Water and mucus samples were frozen immediately after collection on dry ice and were subjected to shotgun metagenomic sequencing.

### Sequencing

DNA purifications from the carp skin mucus samples were performed in triplicates and the resulting total DNA extracts were pooled together. All extractions were carried out using ZymoBIOMICS DNA/RNA miniprep kits (R2002, Zymo Research, Irvine, USA). For efficient lysis of the mucus samples, bead homogenization was performed using a Vortex-Genie 2 with a bead size of 0.1 mm, a homogenization time of 15 min at maximum speed, after which the Zymo Research kit’s DNA purification protocol was followed. Total DNA qualities were assessed with an Agilent 2200 TapeStation instrument (Agilent Technologies, Santa Clara, USA), and DNA quantities were measured using a Qubit Flex Fluorometer.

We closely followed all manufacturer recommendations when preparing sequencing libraries for Illumina sequencing platform (Illumina Inc., San Diego, USA). Pooled total DNA samples were used to construct libraries using the NEBNext Ultra II Library Prep Kit (NEB, Ipswich, MA, USA). Paired-end shotgun metagenome sequencing was performed on a NextSeq 550 (Illumina) sequencer using the NextSeq High Output Kit v2 sequencing reagent kit. Primary data analysis (i.e., base-calling) was performed using “bcl2fastq” software (version 2.17.1.14, Illumina).

### Bioinformatic Analysis

Quality control of the raw reads was performed with FastQC v0.11.9^29^ and MultiQC v1.11^30^. Read pairs were merged with PEAR v0.9.11^31^ before quality filtering. However, to retain as much information as possible for downstream analysis forward reads of those pairs that couldn’t be merged were also used. TrimGalore v0.6.7^32^ was used for quality trimming of the merged and forward unmerged (see above) reads. After these steps, reads were dereplicated with VSEARCH v2.18.0^33^. Taxonomic classification of the reads was performed with Kraken v2.1.2^34^ to the NCBI nt database (built on: 26.12.2022).

Following classification, the bacteriome of samples was analyzed in R v4.1.2 environment^35^. The taxonomic composition of samples was examined with the aid of the phyloseq v1.38.0^36^ R Bioconductor package. Rarefaction curves were calculated with the vegan v2.6-2^37^ package at the species level. Good’s coverage estimator was adapted based on Lin et al.^38^ at the species level by the following formula: (1 − (*n/N*)) * 100, where *n* is the number of species with only one classified read, and *N* is the number of all reads classified at the species level.

Shannon index^39^ was calculated with the phyloseq package to infer alpha-diversity of samples at species level classification. To avoid bias caused by the unequal sequencing depth between samples, Shannon indices were calculated as an average of 1000 iterations of random rarefications of each sample to an even depth (number of reads after rarefication: 6274). Alongside the mean Shannon indices from the random rarefications, the average number of observed species was also calculated.

Furthermore, bacteriome was analyzed at the phylum level composition. However, as it can be important for a broad understanding of the bacterial community at hand and might be beneficial when comparing different hosts, deeper knowledge and inferences on its function can only be made by analyzing the structure at finer taxonomic levels as well. We aimed to obtain a comprehensive description of the bacterial composition of the common carp skin mucus and consequently decided to analyze all of the dominant phyla at the genus level in more detail. We believe that a more even description can be provided by this method on the bacterial genera present in the external mucus of the carp than would be possible by a global threshold on genus-level relative abundances. Dominant phyla were selected if their relative abundance had reached at least 1% in any of the fish samples analyzed.

For functional prediction of the bacteriome, reads classified as originating from bacteria were assembled to contigs by MEGAHIT v1.2.9^40^. Genes on the resulting contigs were predicted and converted to protein sequences with Prodigal v2.6.3^41^. Functional annotation of these sequences was performed with eggNOG-mapper v2.1.10^42,43^ to the eggNOG bacterial HMM database^44^ (built on: 01.03.2023). For the carp skin mucus samples, annotated genes were pooled for samples originating from the same pond for the functional analysis. Redundant Gene Ontology (GO) Biological Process terms^45,46^ associated with the annotated genes were filtered with REVIGO^47^ at dispensability cut off of 0.5. Visualization of GO Biological Process terms was performed by multidimensional scaling of the terms based on their semantic similarity as performed by REVIGO. Furthermore, KEGG (Kyoto Encyclopedia of Genes and Genomes)^48–50^ pathway enrichment analysis was performed on the annotated genes with the enrichKEGG function of the ClusterProfiler^51^ R Bioconductor package with Benjamini-Hochberg corrected p-value cutoff of 0.05.

## Results

### Skin mucus bacterial community

Due to the high contamination by the host genome in our fish skin mucus samples (the percentage of the kingdom Bacteria (mean *±* SD) was 0.12 *±* 0.12 in fish skin mucus, whereas it was 70.65 *±* 0.47 in water samples) the rarefaction curves did not reach their plateau. Even though this might limit our conclusions on the bacteriome composition of the common carp skin mucus, our samples still provide valuable insight into the main constitution of fish skin mucus bacteriome. Rarefaction curves are presented in Supplementary Figure 1, while Good’s coverage values can be found in Supplementary File 2.

Shannon index was calculated for each sample (either skin mucus or water) to obtain a broader understanding of the communities by their alpha-diversity. Shannon index values and the number of observed species are summarized for each sample in 1.

The bacterial composition of samples was analyzed at different taxonomic levels to gain a deeper understanding on the skin bacteriome of carp. Dominant bacterial phyla and their relative abundance are presented in 1 for each carp skin sample and the two water samples as well. A phylum was considered dominant if its abundance had reached at least 1% in any of the samples analyzed. According to this threshold *Proteobacteria, Actinobacteria, Bacteroidota, Firmicutes, Cyanobacteria* and *Planctomycetota* were regarded as dominant and was selected for further genus level description.

Relative abundances of bacterial genera found within these phyla are presented in Supplementary Figures 2-7. For better readability of the plots and to highlight the more substantial genera based on relative abundance, those of them that did not reach 1% relative abundance in at least one sample were aggregated in the artificial category termed “Others”. The distribution of the relative abundance values associated with this category for each bacterial phylum included in the genus level analysis is presented in Supplementary Figure 8. *Planctomycetota* and *Cyanobacteria* have the lowest values for the relative distribution for the category “Others”, while in the case of *Proteobacteria*, even the lowest value is above 40%, indicating the high number of low abundance genera considering that phylum. Relative abundance values used for 1 and Supplementary Figures 2-7 are presented in Supplementary File 3.

**Table 1.** Alpha-diversity measures. Shannon index and the number of observed species were calculated for each fish skin and water sample based on the average of 1000 random rarefications of the shotgun metagenomic reads aligned to the bacteria kingdom and classified at the species level.

| Sample ID | Sample Type | Pond ID | Shannon index | Observed Species count |
| --- | --- | --- | --- | --- |
| SM1 | carp skin mucus | 1 | 6.98 | 2319.71 |
| SM2 | carp skin mucus | 1 | 7.40 | 2677.66 |
| SM3 | carp skin mucus | 1 | 7.19 | 2518.51 |
| SM4 | carp skin mucus | 1 | 7.08 | 2644.11 |
| SM5 | carp skin mucus | 2 | 7.43 | 2731.13 |
| SM6 | carp skin mucus | 2 | 7.49 | 2801.81 |
| SM7 | carp skin mucus | 2 | 6.80 | 2106.29 |
| SM8 | carp skin mucus | 2 | 7.29 | 2485.66 |
| W1 | water | 1 | 7.32 | 2707.35 |
| W2 | water | 2 | 7.35 | 2668.29 |

**Figure 1.**
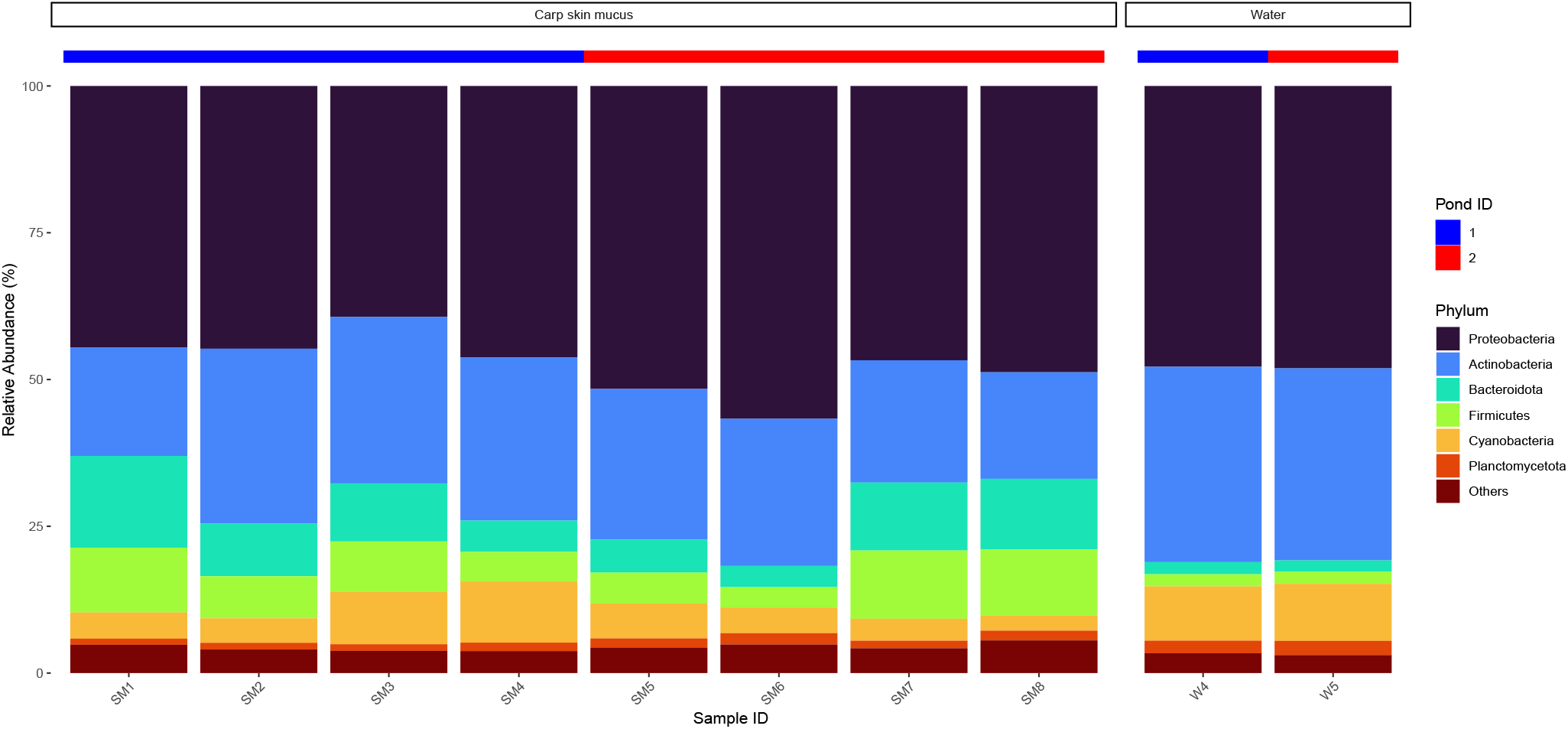
Relative abundance of the dominant bacterial phyla in each sample. Samples are presented on the x-axis, and corresponding bars show the relative abundance of phyla found in our analysis. Water and carp skin mucus samples are separated, and the ID of the pond the samples are originating from is indicated above the bars. Phyla that did not reach at least 1% in at least 1 sample were aggregated in the artificial category termed “Others”.

### Functional prediction of the skin mucus bacteriome

Biological functions associated with the skin bacteriome of the carp were inferred from the genes annotated by eggNOG-mapper. Enrichment of KEGG pathways among the pooled genes for each pond was performed with the ClusterProfiler R Bioconductor package, however, in the case of pond 1, only the gene set associated with quorum sensing was found to be significant (Benjamini-Hochber corrected p = 0.0038). KEGGpathway enrichment analysis of the genes from pond 2 is presented in 2 **A**. Considering the number of Gene Ontology Biological Process terms associated with genes found on carp skin mucus samples from each pond were 211 and 443 for ponds 1 and 2, respectively. The remaining GO terms after redundancy filtering by REVIGO can be found in 2 **B** and **C**. Supplementary Figures 9-10 show the KEGG pathway enrichment results for the water samples. All significantly enriched KEGG pathways and GO terms associated with the annotated genes for the skin mucus and water samples from each pond can be found in Supplementary File 4.

## Discussion

The bacteriome of the fish external mucus has an important function for the health of its host as a primary line of defence^3–5^. Despite its importance, only a few studies have focused on the composition of the skin bacteriome of fishes, with the majority being 16S rRNA gene-based investigations^4,14,21^, which can only provide limited information on its function compared to shotgun metagenomic approaches^14,24,25^. For common carp, which is of major economic importance^16–18^, there is even less information available on its microbiome, especially considering the skin mucus. For these reasons, our results may provide important insight into the composition of the bacteriome colonizing the outer surfaces of common carp and the various functions associated with the genes it encodes, which could serve as a basis for future research.

We have analyzed the bacteriome composition of our samples from different aspects. First, we compared the alpha-diversity of samples, a measure that can be used to summarise the composition of populations according to the number of taxa they contain and their distribution^52,53^. In addition to being an important ecological characteristic of a community, alpha-diversity can also have an important practical role. In human cases, it has been described as an indicator of several disease states^54,55^, and may even be associated with response to therapy in the treatment of certain conditions^56^. Disease has often been associated with a decrease in alpha diversity, but the opposite has also been reported^57^. It is important to note that, in addition to humans, alpha diversity has also been shown to be associated with various diseases in fish as well^58,59^.

The Shannon diversity index^39^ was used to summarise alpha-diversity in our samples, which is one of the most popular measures available^60^. The mean Shannon diversity index calculated at the species level was 7.21 *±* 0.24 (mean *±* SD) in our fish skin mucus samples. Interestingly, the majority of previous studies on the microbiome of the fish external mucus observed lower Shannon index values. In cases with the highest Shannon diversity, index measurements around 5 were observed, but the majority had even lower values^7,61–63^. An exception to the above is the study by Mohammed and Arias^5^, where the effect of potassium permanganate on *Flavobacterium columnare* in channel catfish was investigated. Here, the Shannon diversity of skin and gill samples from the pre-treatment and control groups was 7.31 and 7.60, respectively.

**Figure 2.**
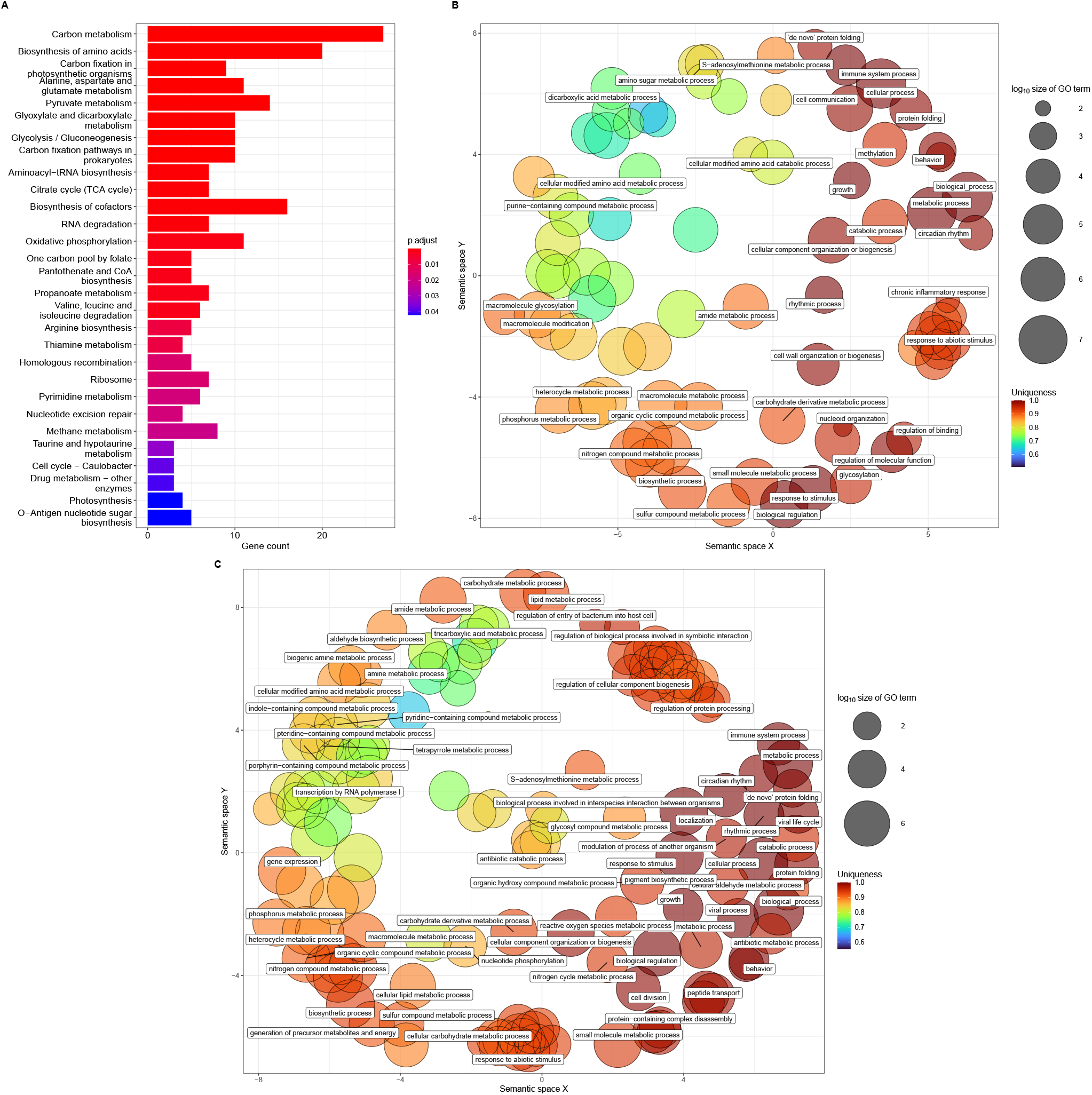
KEGG enrichment and GO terms associated with genes annotated for carp skin samples. KEGG enrichment result of the pooled genes annotated by eggNOG-mapper from carp skin samples from pond 2 are shown in **A**. Bar size along the x-axis indicates the number of genes found in our analysis corresponding to that pathway. Colors indicate the Benjamini-Hochberg corrected p-values. Gene Ontology Biological Process terms are presented after redundancy filtering by REVIGO^47^ for carp skin samples from ponds 1 and 2 at **B** and **C**, respectively. Biological process terms are positioned by multidimensional scaling, considering their similarity. The size of circles indicates the log_10_ transformed number of terms annotated for the corresponding Gene Ontology term. Uniqueness, as calculated by REVIGO, indicates the similarity of terms to others supplied for redundancy filtering. Circles are colored according to the uniqueness of the corresponding term. For better readability, only those terms are written to the plot which has dispensability values associated with them lower than 0.2.

Although the difference in diversity measures may be due to several reasons, one of the major causes could be the difference between fish species and the environmental conditions that samples originate from. Furthermore, the importance of the different technical factors should be noted as well. An important factor could be the data used for the calculation of the Shannon index. For example, Minniti et al.^61^, and Mohammed and Arias^5^, used the OTU level classification for calculation, however in most cases, it was not specified^7,62,63^. In addition, in the case of 16S rRNA gene-based approaches, it is generally considered that they yield poorer resolution compared to shotgun metagenomics^26,27^, which can bias estimates. It is worth noting that the difference in Shannon diversity between our results and previous studies was observed despite the failure to saturate rarefaction curves in our carp samples due to high levels of host genome contamination.

It would have been useful to compare our results with other *C. carpio* skin mucus samples, but to the best of our knowledge, such analyses have not yet been performed, even with 16S rRNA gene sequencing. Furthermore, we also were not able to make comparisons with shotgun metagenome sequencing-based studies on other species because the study on eel skin metagenome from Carda-Diéguez et al.^22^ did not specify the method used for calculating alpha-diversity, while in the case of Doane et al.^23^, no alpha-diversity was calculated from the samples.

However, by examining the bacteriome in detail, we can obtain much more information about its composition and function than diversity alone can tell us. Based on the taxonomic constitution of our samples, *Proteobacteria* and *Actinobacteria* phyla were clearly dominant both in fish skin mucus and water samples. The dominance of the *Proteobacteria* phylum is not an uncommon observation in fish external mucus samples^1,3,5,6,8,11,21,62,63^, however, differences between fish species have been observed for the other phyla^1,11,62,63^. Moreover, significant within-species variability in dominant phyla has been described^64^, and variability within individuals related to body sites should be noted^12^.

The microbiome can be an important indicator of various pathological conditions, which has already been described in fish, for example, in the case of the gastrointestinal tract^65^. In this regard, the *Bacteroidota* phylum may be interesting, which has been highlighted as a marker for eutrophication^9,66^. Understanding the changes in the composition of the bacteriome or even the microbiome during different pathological conditions can be an important step in understanding and potentially diagnosing disease processes.

Our results are therefore in line with the dominance of the *Proteobacteria* phylum observed in other fish species, but direct comparison with *C. carpio* is not possible due to the lack of available data. Of course, our observations on the bacteriome composition of our samples are also limited by their paramount host genome contamination, which reduced the coverage of bacterial genomes of interest in the sequencing reaction.

Shotgun sequencing allows us to functionally analyze samples using the genes annotated from reads aligned to bacterial genomes^14,24,25^. Genes annotated with eggNOG-mapper are also assigned with Gene Ontology Biological Functions by the database. These, together with pathways found to be significant by KEGG enrichment analysis, revealed various metabolic processes in the carp mucus samples. The outer mucus of fish provides a nutrient-rich environment for microorganisms^3,21^ to metabolize, however, KEGG enrichment analysis also found metabolic pathways related to photosynthesis as well. A significant fraction of the mucin lining of fish skin is composed of glycosylated proteins^67^, so it is not surprising that the GO function “glycosyl compound metabolic process” could have been linked to one or more annotated genes.

In the GO Biological Function set, we also had some interesting hits, such as “regulation of entry of bacterium into host cell” and “biological process involved in interspecies interaction between organisms,” which may be related to the relationship between the bacteriome and the host. Furthermore, ontologies “chronic inflammation” and “immune system process” may indicate that even host-derived genome sequences may have appeared in the assemblies, even though they were only calculated from the reads aligning to the Bacteria kingdom. In addition to this, the “antibiotic catabolic process” function can be highlighted as it may be related to the resident bacteriome trying to prevent the colonization of other bacteria^5,21^, although we cannot exclude the possibility of external antibiotic pressure.

The skin of fish is in direct contact with the surrounding water, and several studies have compared the bacteriome of the external mucus to the surrounding water. Most of the results to date have found that the two communities have different bacterial assemblages^8,12,21,64^, however, the opposite has also been reported^9^. In our case, we found a very high degree of similarity at the phylum level and also at the genus level within the dominant phyla. This is well illustrated by the genera corresponding to the *Cyanobacteria* phylum, where the difference between the water from two ponds in the proportion of genera *Plankthothrix* and *Synechoccus* is well reflected in the carp samples from the respective ponds (see Supplementary File 2 for more details). Beyond this, of course, there were genera that showed differences between water and carp samples, such as the *Delftia* genus being more abundant in the fish slime, while the *Methylocystis* genus was more pronounced in the water samples in the case of the *Proteobacteria* phylum (for more details, please see Supplementary File). Our results, therefore, support the similarity of the bacterial composition of the fish mucus and the surrounding water, at least for *C. carpio*. It is important to note, however, that comparisons of skin mucus and water samples from different species and different environments are not straightforward and do not necessarily give generally valid results. However, a deeper understanding of the relationship between the two environments raises interesting further questions.

## Conclusion

To the best of our knowledge, our results are the first to describe the bacterial community of the mucus that covers the skin of *C. carpio*. Considering the economic importance of this species and the important role that the microbiome can play in understanding and detecting various pathological conditions, studies of the bacteriome could provide valuable data. Our results have demonstrated that shotgun metagenomics can provide useful information on the skin bacteriome of this species, with particular emphasis on the potential for functional studies. A particularly important observation of our analysis is that the host genome can significantly contaminate the samples, and even though it still can provide useful information, it is preferable to be prepared in advance when designing experiments. Furthermore, our results may provide a useful basis for comparison for future studies in common carp.

## Supplementary Information

**Supplementary File 1:** Metadata and read counts for each sample. Each row in the table represents one sample. Columns correspond to the sample ID (**Sample ID**) used in the present study, sample type (i.e. the origin of the samples, **Sample Type**), the phenotype of the specimen (**Phenotype**), the presence of ulcer on the skin of the specimen (**Ulcer**), the unique ID of the pond the sample is originating from (**Pond ID**), read counts in the specific samples after each step of the bioinformatic analysis (**Number of read pairs, Number of merged reads, Number of trimmed merged reads**), the accuracy of the Kraken2 taxonomic classification (i.e. the ratio of the reads classified, **Classification accuracy (%)**) and the Good’s coverage estimator values for each sample (**Good’s coverage**).

**Supplementary File 2:** Supplementary PDF file containing Supplementary Figures 1-10.

**Supplementary File 3:** Excel file of the relative abundance values for each bacterial phyla and bacterial genera amongst their respective phyla for *Proteobacteria, Actinobacteria, Bacteroidota, Firmicutes, Cyanobacteria* and *Planctomycetota*. Data for the phylum level abundance values and the genus level abundances for each phylum are presented on separate sheets in the Excel file. Columns of the tables correspond to the sample IDs (**Sample ID**), type of the samples (**Sample Type**), the ID of the pond (**Pond ID**), the relative abundance of the given phylum/genus in the corresponding sample (**Relative abundance**) and the name of the phylum/genus (**Phylum** or **Genus**). For the genus level analysis, those genera that did not reach at least 1% relative abundance in at least one sample were aggregated to the artificial category “Others”.

**Supplementary File 4:** Excel file containing the results of KEGG enrichment analyses and Gene Ontology Biological Process terms after redundancy filtering by REVIGO. The first four Excel sheets show the KEGG enrichment results for the pooled genes from carp skin mucus from ponds 1 and 2 and water samples from ponds 1 and 2. Columns correspond to the KEGG ID of the pathway found to be significant (**KEGG ID**), the description of the pathway (**Description**), p-value (column **P-value**) and Benjamini-Hochberg adjusted p-value (**Adjusted P-value**), KEGG Ontology IDs of the genes found to be associated with the corresponding pathway (**KEGG Ontology ID**) and the number of these genes (**Gene Count**). The last two sheets contain the Gene Ontology Biological Process terms after redundancy filtering with REVIGO. Columns correspond to the Biological Process term found to be associated with any of the genes found for the carp skin mucus samples for the corresponding pond (**GO BP Term ID**), the name of the term (**Name**), log_10_ size of the term (**Log10 Size**), frequency of the term in the database (i.e. the generality of the term, column **Frequency**), uniqueness (column **Uniqueness**) and dispensability (column **Dispensability**) as calculated by REVIGO. The last column (**Representative Term**) indicates the Gene Ontology term ID of the cluster representative that the term was assigned to by REVIGO, if the column contains the value “null”, that means that the term is the representative itself.

## Declarations

### Ethics approval and consent to participate

Not applicable.

### Consent for publication

Not applicable.

### Availability of data and material

Sequencing data is publicly available and can be accessed through the PRJNA982406 BioProject from the Sequence Read Archive (SRA) at NCBI.

### Author contributions statement

MP and NS take responsibility for the data’s integrity and the data analysis’s accuracy. MP and NS conceived the concept of the study. MP, NS, AGT, KE, LC, and MK participated in the sample collection. GM performed the sequencing. MP and NS participated in the bioinformatic analysis and in the drafting of the manuscript. GM, LB, KE, MP, NS, AGT, SÁN, and NC carried out the critical revision of the manuscript for important intellectual content. All authors read and approved the final manuscript.

### Authors’ information

No relevant information can be provided of the authors that may aid the readers’ interpretation of the article.

## References

1. Boutin, S., Bernatchez, L., Audet, C. & Derôme, N. Network analysis highlights complex interactions between pathogen, host and commensal microbiota. PloS ONE 8, e84772 (2013).

2. Guerrero, R., Margulis, L., Berlanga, M. et al. Symbiogenesis: the holobiont as a unit of evolution. Int. Microbiol. 16, 133–143 (2013).

3. Merrifield, D. L. & Rodiles, A. The fish microbiome and its interactions with mucosal tissues. In Mucosal health in aquaculture, 273–295 (Elsevier, 2015).

4. Llewellyn, M. S., Boutin, S., Hoseinifar, S. H. & Derome, N. Teleost microbiomes: the state of the art in their characterization, manipulation and importance in aquaculture and fisheries. Front. Microbiol. 5, 207 (2014).

5. Mohammed, H. H. & Arias, C. R. Potassium permanganate elicits a shift of the external fish microbiome and increases host susceptibility to columnaris disease. Vet. Res. 46, 82 (2015).

6. Sylvain, F.-É. et al. pH drop impacts differentially skin and gut microbiota of the Amazonian fish tambaqui (Colossoma macropomum). Sci. Reports 6, 1–10 (2016).

7. Xavier, R. et al. A risky business? Habitat and social behavior impact skin and gut microbiomes in Caribbean cleaning gobies. Front. Microbiol. 10, 716 (2019).

8. Sylvain, F.-É. et al. Fish skin and gut microbiomes show contrasting signatures of host species and habitat. Appl. Environ. Microbiol. 86, e00789–20 (2020).

9. Côte, J. et al. Changes in fish skin microbiota along gradients of eutrophication in human-altered rivers. FEMS Microbiol. Ecol. 98, fiac006 (2022).

10. Chiarello, M. et al. Skin microbiome of coral reef fish is highly variable and driven by host phylogeny and diet. Microbiome 6, 1–14 (2018).

11. Larsen, A., Tao, Z., Bullard, S. A. & Arias, C. R. Diversity of the skin microbiota of fishes: evidence for host species specificity. FEMS Microbiol. Ecol. 85, 483–494 (2013).

12. Chiarello, M., Villeger, S., Bouvier, C., Bettarel, Y. & Bouvier, T. High diversity of skin-associated bacterial communities of marine fishes is promoted by their high variability among body parts, individuals and species. FEMS Microbiol. Ecol. 91, fiv061 (2015).

13. Sehnal, L. et al. Microbiome composition and function in aquatic vertebrates: small organisms making big impacts on aquatic animal health. Front. Microbiol. 358 (2021).

14. Legrand, T. P., Wynne, J. W., Weyrich, L. S. & Oxley, A. P. A microbial sea of possibilities: current knowledge and prospects for an improved understanding of the fish microbiome. Rev. Aquac. 12, 1101–1134 (2020).

15. Balcázar, J. L. et al. The role of probiotics in aquaculture. Vet. Microbiol. 114, 173–186 (2006).

16. FAO. The State of World Fisheries and Aquaculture 2022. Towards Blue Transformation. (Food and Agriculture Organization of the United Nations, Rome, 2022).

17. FEAP. Annual Report, 2022 (Federation of European Aquaculture Procedures, 2022).

18. Lasner, T., Mytlewski, A., Nourry, M., Rakowski, M. & Oberle, M. Carp land: Economics of fish farms and the impact of region-marketing in the Aischgrund (DEU) and Barycz Valley (POL). Aquaculture 519, 734731 (2020).

19. Reed, G. B. & Spence, C. M. The intestinal and slime flora of the haddock: a preliminary report. Contributions to Can. Biol. Fish. 4, 257–264 (1929).

20. Horsley, R. A review of the bacterial flora of teleosts and elasmobranchs, including methods for its analysis. J. Fish biology 10, 529–553 (1977).

21. Gomez, J. A. & Primm, T. P. A slimy business: the future of fish skin microbiome studies. Microb. Ecol. 1–13 (2021).

22. Carda-Diéguez, M., Ghai, R., Rodríguez-Valera, F. & Amaro, C. Wild eel microbiome reveals that skin mucus of fish could be a natural niche for aquatic mucosal pathogen evolution. Microbiome 5, 1–15 (2017).

23. Doane, M. P. et al. The skin microbiome of elasmobranchs follows phylosymbiosis, but in teleost fishes, the microbiomes converge. Microbiome 8, 1–15 (2020).

24. Jovel, J. et al. Characterization of the gut microbiome using 16S or shotgun metagenomics. Front. Microbiol. 7, 459 (2016).

25. Clooney, A. G. et al. Comparing apples and oranges?: next generation sequencing and its impact on microbiome analysis. PloS ONE 11, e0148028 (2016).

26. Laudadio, I. et al. Quantitative assessment of shotgun metagenomics and 16S rDNA amplicon sequencing in the study of human gut microbiome. OMICS: A J. Integr. Biol. 22, 248–254 (2018).

27. Durazzi, F. et al. Comparison between 16S rRNA and shotgun sequencing data for the taxonomic characterization of the gut microbiota. Sci. Reports 11, 1–10 (2021).

28. Papp, M. et al. Natural diversity of the honey bee (Apis mellifera) gut bacteriome in various climatic and seasonal states. PloS ONE 17, e0273844 (2022).

29. Andrews, S. FastQC: a quality control tool for high throughput sequence data. https://www.bioinformatics.babraham.ac.uk/projects/fastqc/ (2020).

30. Ewels, P., Magnusson, M., Lundin, S. & Käller, M. Multiqc: summarize analysis results for multiple tools and samples in a single report. Bioinformatics 32, 3047–3048 (2016).

31. Zhang, J., Kobert, K., Flouri, T. & Stamatakis, A. PEAR: a fast and accurate Illumina Paired-End reAd mergeR. Bioinformatics 30, 614–620 (2014).

32. Krueger, F. TrimGalore: A wrapper around Cutadapt and FastQC to consistently apply adapter and quality trimming to FastQ files, with extra functionality for RRBS data. https://github.com/FelixKrueger/TrimGalore, DOI: 10.5281/zenodo.7598955 (2021).

33. Rognes, T., Flouri, T., Nichols, B., Quince, C. & Mahé, F. VSEARCH: a versatile open source tool for metagenomics. PeerJ 4, e2584 (2016).

34. Wood, D. E., Lu, J. & Langmead, B. Improved metagenomic analysis with Kraken 2. Genome biology 20, 1–13 (2019).

35. R Core Team. R: A Language and Environment for Statistical Computing. R Foundation for Statistical Computing, Vienna, Austria (2021).

36. McMurdie, P. J. & Holmes, S. phyloseq: An R package for reproducible interactive analysis and graphics of microbiome census data. PloS ONE 8, e61217 (2013).

37. Oksanen, J. et al. vegan: Community Ecology Package (2022). R package version 2. 6-2.

38. Lin, A. et al. Distinct distal gut microbiome diversity and composition in healthy children from Bangladesh and the United States. PloS ONE 8, e53838 (2013).

39. Shannon, C. E. A mathematical theory of communication. The Bell system technical journal 27, 379–423 (1948).

40. Li, D., Liu, C.-M., Luo, R., Sadakane, K. & Lam, T.-W. MEGAHIT: an ultra-fast single-node solution for large and complex metagenomics assembly via succinct de Bruijn graph. Bioinformatics 31, 1674–1676 (2015).

41. Hyatt, D. et al. Prodigal: prokaryotic gene recognition and translation initiation site identification. BMC Bioinform. 11, 1–11 (2010).

42. Eddy, S. R. Accelerated profile HMM searches. PLoS Comput. Biol. 7, e1002195 (2011).

43. Cantalapiedra, C. P., Hernández-Plaza, A., Letunic, I., Bork, P. & Huerta-Cepas, J. eggNOG-mapper v2: functional annotation, orthology assignments, and domain prediction at the metagenomic scale. Mol. Biol. Evol. 38, 5825–5829 (2021).

44. Huerta-Cepas, J. et al. eggNOG 5.0: a hierarchical, functionally and phylogenetically annotated orthology resource based on 5090 organisms and 2502 viruses. Nucleic Acids Res. 47, D309–D314 (2019).

45. Ashburner, M. et al. Gene ontology: tool for the unification of biology. Nat. genetics 25, 25–29 (2000).

46. The Gene Ontology resource: enriching a GOld mine. Nucleic Acids Res. 49, D325–D334 (2021).

47. Supek, F., Bošnjak, M., Škunca, N. & Šmuc, T. REVIGO summarizes and visualizes long lists of gene ontology terms. PloS ONE 6, e21800 (2011).

48. Kanehisa, M. & Goto, S. KEGG: kyoto encyclopedia of genes and genomes. Nucleic Acids Res. 28, 27–30 (2000).

49. Kanehisa, M. Toward understanding the origin and evolution of cellular organisms. Protein Sci. 28, 1947–1951 (2019).

50. Kanehisa, M., Furumichi, M., Sato, Y., Kawashima, M. & Ishiguro-Watanabe, M. KEGG for taxonomy-based analysis of pathways and genomes. Nucleic Acids Res. 51, D587–D592 (2023).

51. Yu, G., Wang, L.-G., Han, Y. & He, Q.-Y. clusterProfiler: an R package for comparing biological themes among gene clusters. OMICS: A J. Integr. Biol. 16, 284–287 (2012).

52. Magurran, A. E. Measuring biological diversity. Curr. Biol. 31, R1174–R1177 (2021).

53. Willis, A. D. Rarefaction, alpha diversity, and statistics. Front. Microbiol. 10, 2407 (2019).

54. Gong, D., Gong, X., Wang, L., Yu, X. & Dong, Q. Involvement of reduced microbial diversity in inflammatory bowel disease. Gastroenterol. Res. Pract. 2016 (2016).

55. Manor, O. et al. Health and disease markers correlate with gut microbiome composition across thousands of people. Nat. Commun. 11, 5206 (2020).

56. Gopalakrishnan, V., Helmink, B. A., Spencer, C. N., Reuben, A. & Wargo, J. A. The influence of the gut microbiome on cancer, immunity, and cancer immunotherapy. Cancer Cell 33, 570–580 (2018).

57. Avalos-Fernandez, M. et al. The respiratory microbiota alpha-diversity in chronic lung diseases: first systematic review and meta-analysis. Respir. Res. 23, 214 (2022).

58. Mougin, J. & Joyce, A. Fish disease prevention via microbial dysbiosis-associated biomarkers in aquaculture. Rev. Aquac. 15, 579–594 (2023).

59. Li, T. et al. Bacterial signatures of “Red-Operculum” disease in the gut of crucian carp (Carassius auratus). Microb. Ecol. 74, 510–521 (2017).

60. Alberdi, A. & Gilbert, M. T. P. A guide to the application of Hill numbers to DNA-based diversity analyses. Mol. ecology resources 19, 804–817 (2019).

61. Minniti, G. et al. The skin-mucus microbial community of farmed Atlantic salmon (Salmo salar). Front. Microbiol. 8, 2043 (2017).

62. Colin, Y. et al. Urbanization constrains skin bacterial phylogenetic diversity in wild fish populations and correlates with the proliferation of aeromonads. Microb. Ecol. 1–14 (2021).

63. Zhao, N. et al. Heterogeneity of the tissue-specific mucosal microbiome of normal grass carp (Ctenopharyngodon idella). Mar. Biotechnol. 24, 366–379 (2022).

64. Berggren, H. et al. Fish skin microbiomes are highly variable among individuals and populations but not within individuals. Front. Microbiol. 12, 4153 (2022).

65. Luan, Y. et al. The fish microbiota: Research progress and potential applications. Engineering (2023).

66. Krotman, Y., Yergaliyev, T. M., Alexander Shani, R., Avrahami, Y. & Szitenberg, A. Dissecting the factors shaping fish skin microbiomes in a heterogeneous inland water system. Microbiome 8, 1–15 (2020).

67. Reverter, M., Tapissier-Bontemps, N., Lecchini, D., Banaigs, B. & Sasal, P. Biological and ecological roles of external fish mucus: a review. Fishes 3, 41 (2018).

