## Supplementary File 2 for "Shotgun metagenomic analysis of the skin mucus bacteriome of the common carp (*Cyprinus carpio*)"

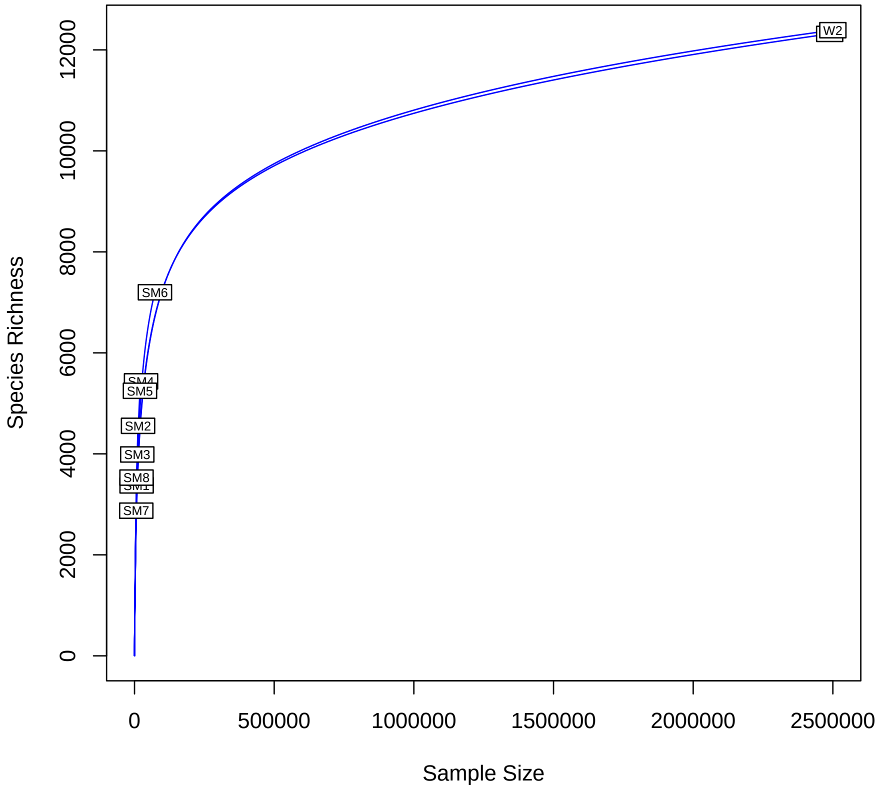


Supplementary Figure 1: Rarefaction curve at the species level of each samples. The curve was calculated with the rarecurve function of the vegan R Bioconductor package.

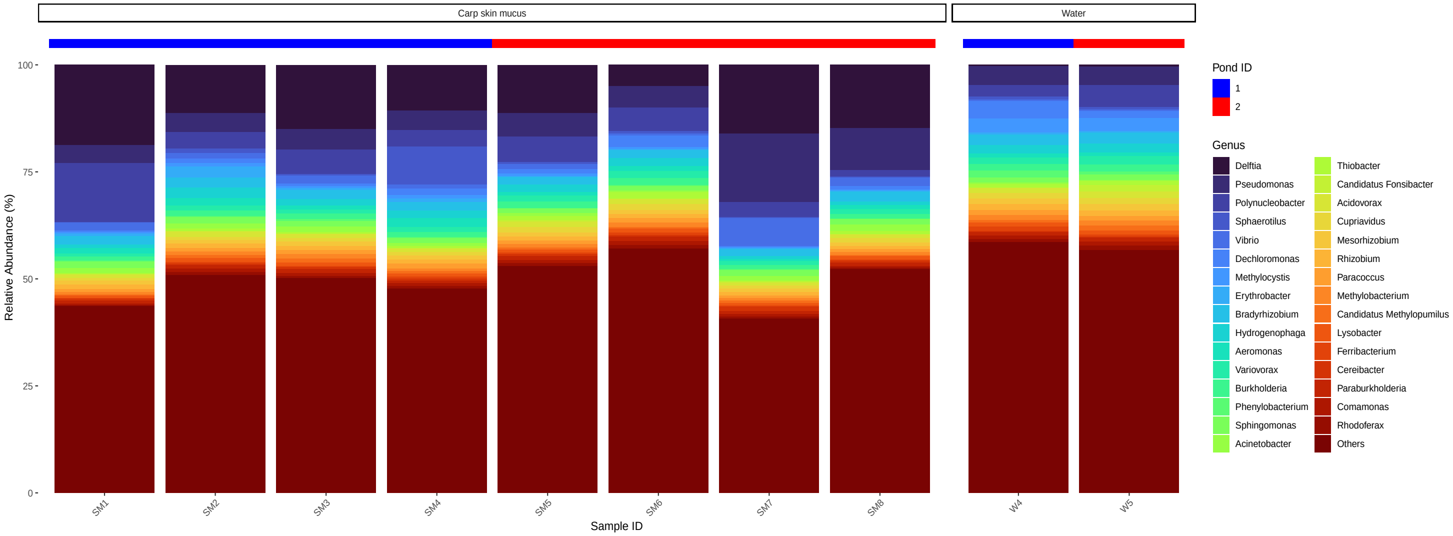


Supplementary Figure 2: Genus level composition of the reads aligned to the *Proteobacteria* phylum for each samples. Genera not reaching at least 1% relative abundance in at least 1 carp skin sample was aggregated in the cathegory Others. Water and carp skin mucus samples are separated and the ID of the pond the samples are originating from is indicated above the bars.

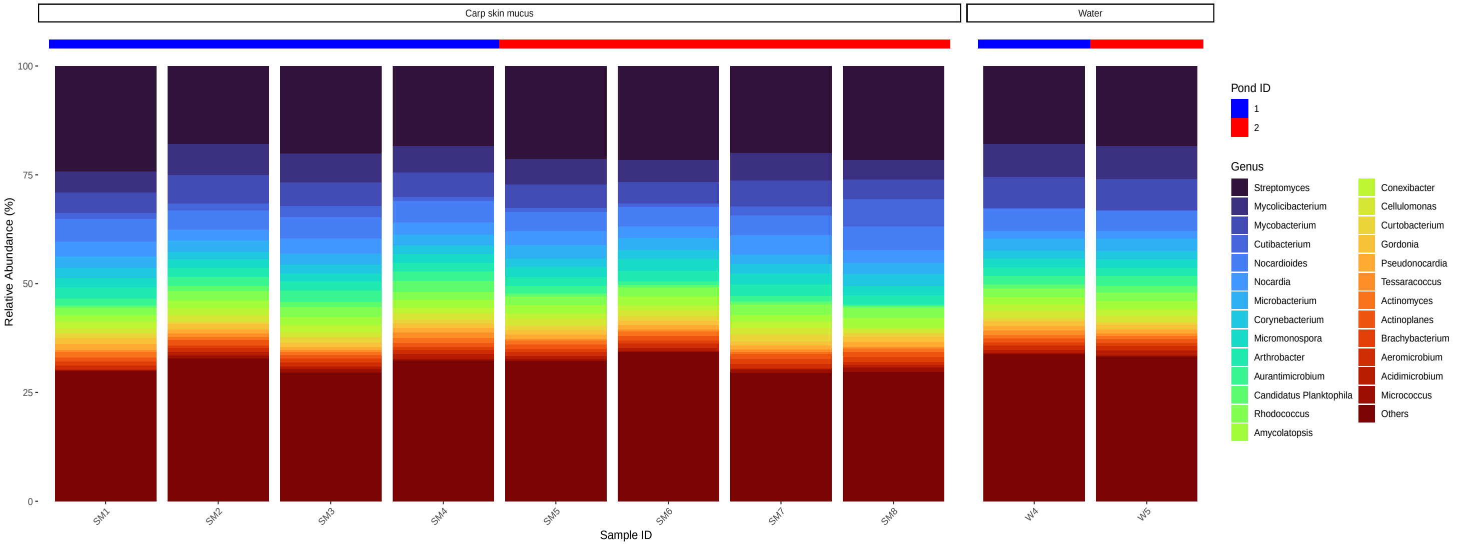


Supplementary Figure 3: Genus level composition of the reads aligned to the *Actinobacteria* phylum for each samples. Genera not reaching at least 1% relative abundance in at least 1 carp skin sample was aggregated in the cathegory Others. Water and carp skin mucus samples are separated and the ID of the pond the samples are originating from is indicated above the bars.

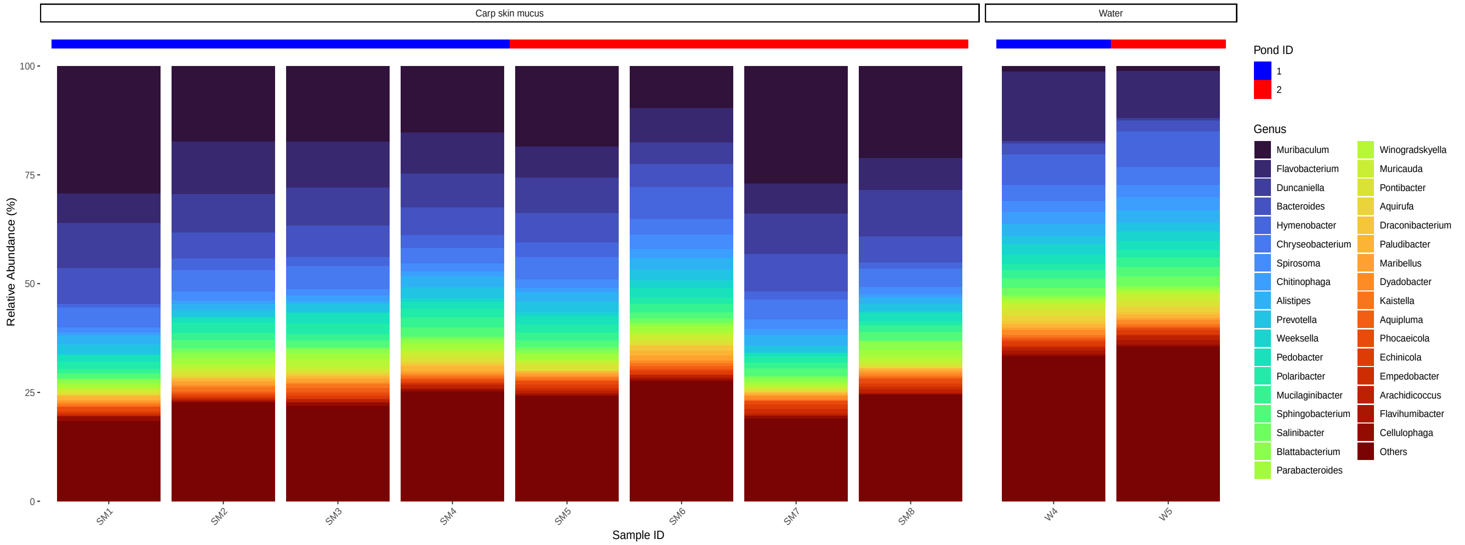


Supplementary Figure 4: Genus level composition of the reads aligned to the *Bacteroidota* phylum for each samples. Genera not reaching at least 1% relative abundance in at least 1 carp skin sample was aggregated in the cathegory Others. Water and carp skin mucus samples are separated and the ID of the pond the samples are originating from is indicated above the bars.

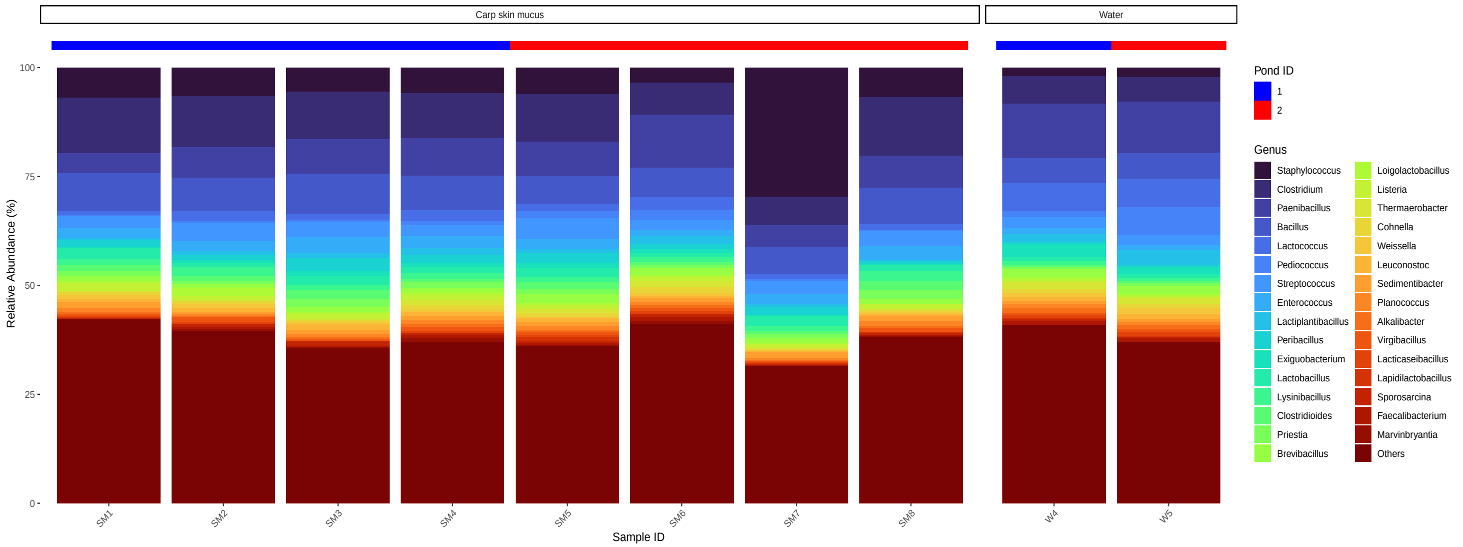


Supplementary Figure 5: Genus level composition of the reads aligned to the *Firmicutes* phylum for each samples. Genera not reaching at least 1% relative abundance in at least 1 carp skin sample was aggregated in the cathegory Others. Water and carp skin mucus samples are separated and the ID of the pond the samples are originating from is indicated above the bars.

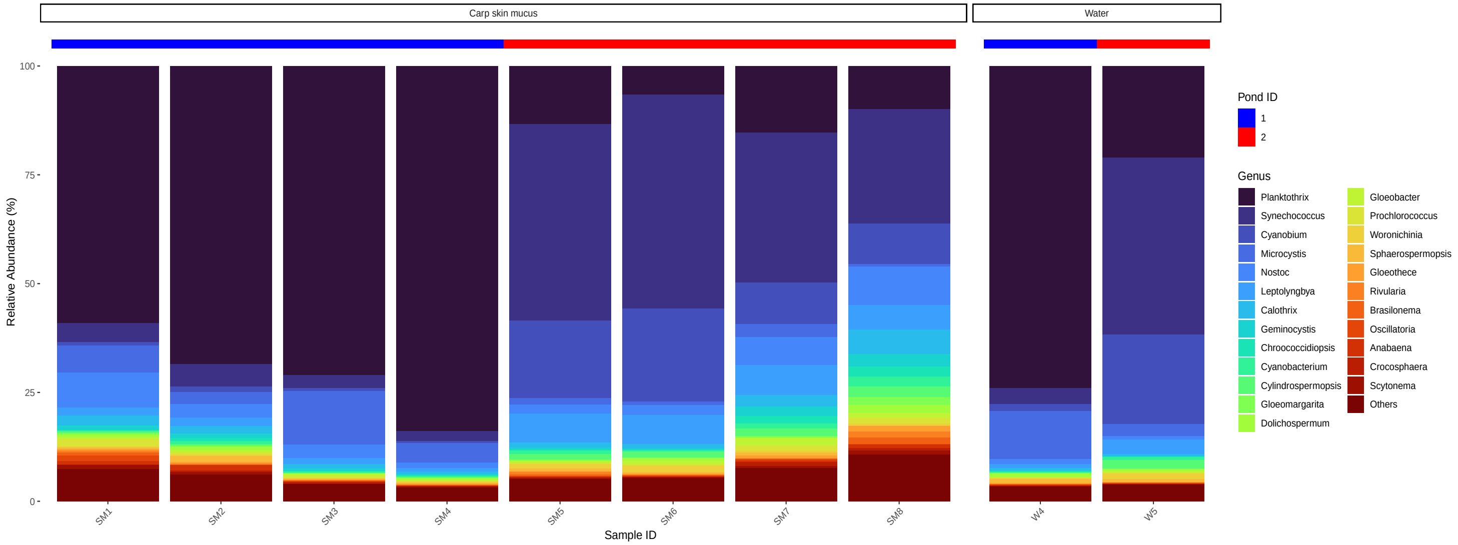


Supplementary Figure 6: Genus level composition of the reads aligned to the *Cyanobacteria* phylum for each samples. Genera not reaching at least 1% relative abundance in at least 1 carp skin sample was aggregated in the cathegory Others. Water and carp skin mucus samples are separated and the ID of the pond the samples are originating from is indicated above the bars.

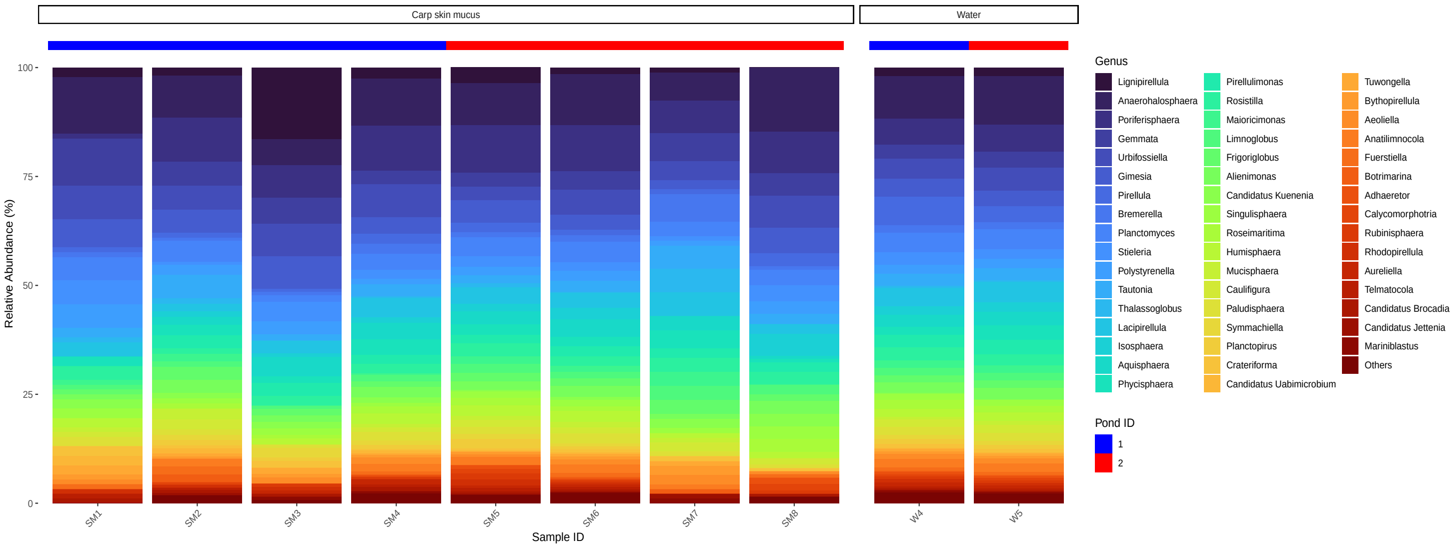


Supplementary Figure 7: Genus level composition of the reads aligned to the *Planctomycetota* phylum for each samples. Genera not reaching at least 1% relative abundance in at least 1 carp skin sample was aggregated in the cathegory Others. Water and carp skin mucus samples are separated and the ID of the pond the samples are originating from is indicated above the bars.

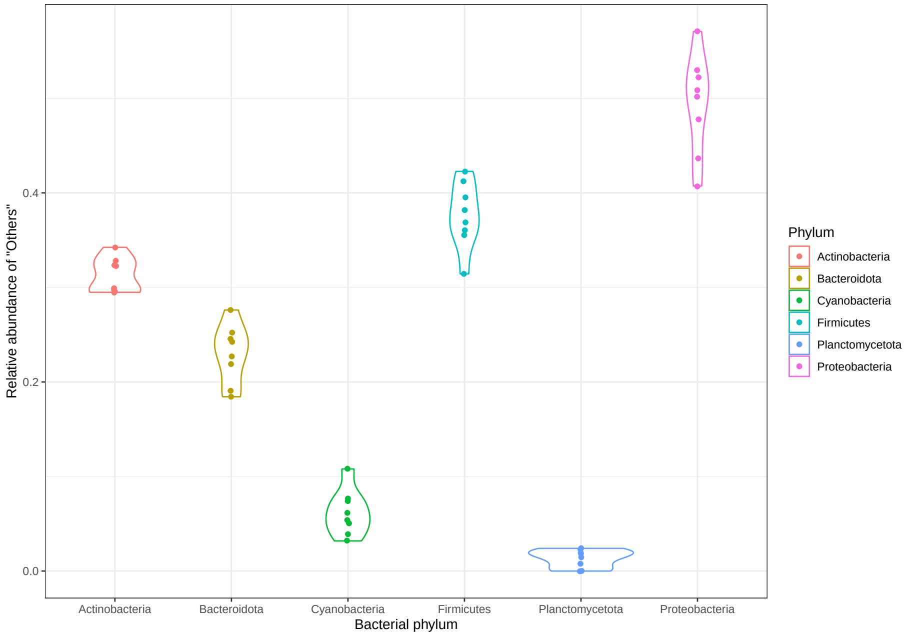


Supplementary Figure 8: Distribution of the relative abundances associated with the cathegory "Others" for each phyla analysed at the genus level in the carp skin mucus samples. The cathegory "Others" was constructed as an aggregate of those genera that did not reach at least 1% relative abundance in at least one sample. Points are jittered along the x axis for better readability of the plot.

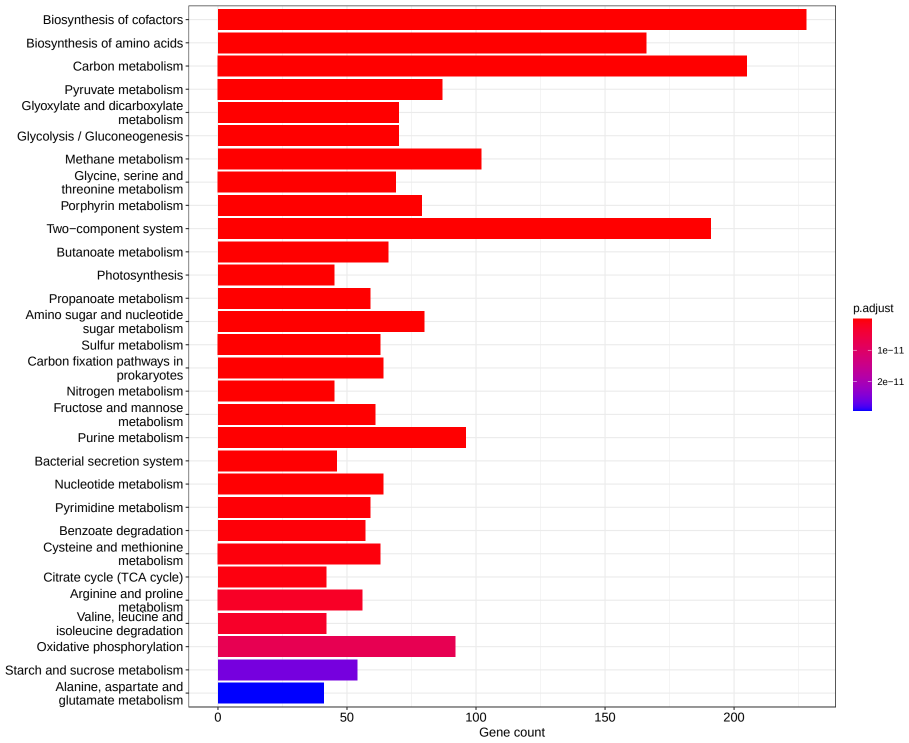


Supplementary Figure 9: KEGG enrichment results of the genes annotated from the pond 1 water sample. The length of bars along the x axis indicate the number of genes found within the given pathway. Colors indicate the Benjamini-Hochberg corrected p-values. The plot was restricted to only the top 30 pathways. For the list of all pathways found significantly enriched in pond 1 water sample see Supplementary File 4.

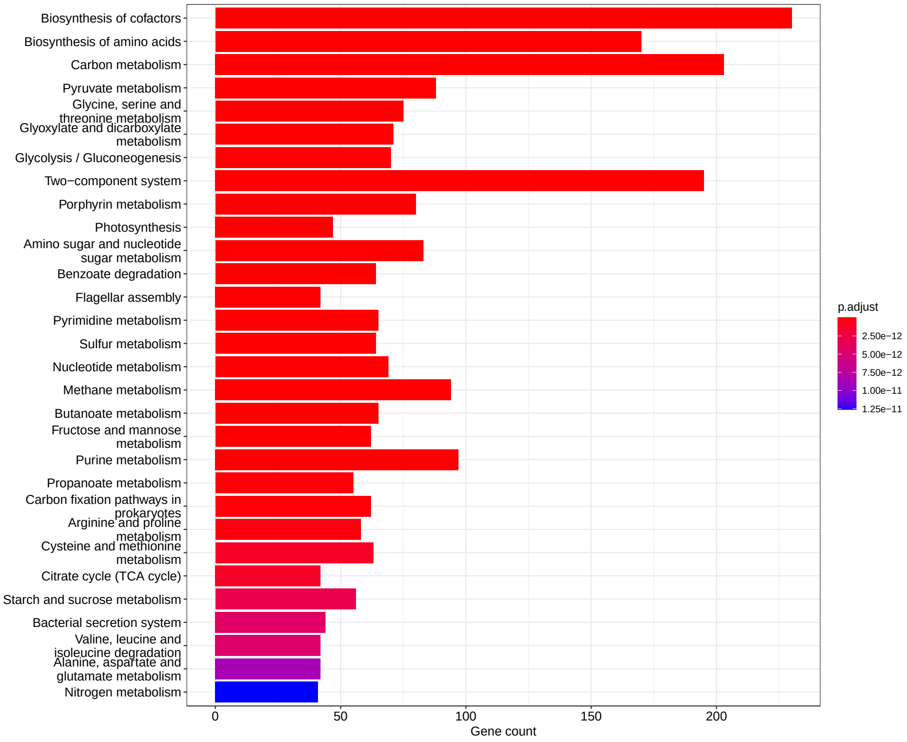


Supplementary Figure 10: KEGG enrichment results of the genes annotated from the pond 2 water sample. The length of bars along the x axis indicate the number of genes found within the given pathway. Colors indicate the Benjamini-Hochberg corrected p-values. The plot was restricted to only the top 30 pathways. For the list of all pathways found significantly enriched in pond 2 water sample see Supplementary File 4.
